# Engineering adoptive T cell therapy to co-opt Fas ligand-mediated death signaling in ovarian cancer enhances therapeutic efficacy

**DOI:** 10.1101/2021.07.30.454539

**Authors:** Kristin G. Anderson, Shannon K. Oda, Breanna M. Bates, Madison G. Burnett, Magdalia Rodgers Suarez, Susan L. Ruskin, Philip D. Greenberg

## Abstract

**Background:** In the U.S., more than 50% of ovarian cancer patients die within 5 years of diagnosis, highlighting the need for innovations such as engineered T cell therapies. Mesothelin (Msln) is an attractive immunotherapy target for this cancer, as it is overexpressed by the tumor and contributes to malignant and invasive phenotypes, making antigen loss disadvantageous to the tumor. We previously showed that adoptively transferred T cells engineered to be Msln-specific (TCR_1045_) preferentially accumulate within established ovarian tumors, delay tumor growth and significantly prolong survival in the ID8_VEGF_ mouse model. However, T cell persistence and anti-tumor activity were not sustained, and we and others have previously detected FasL in the tumor vasculature and the tumor microenvironment (TME) of human and murine ovarian cancers, which can induce apoptosis in infiltrating lymphocytes expressing Fas receptor (Fas).

**Methods:** To concurrently overcome this mechanism for potential immune evasion and enhance T cell responses, we generated an immunomodulatory fusion protein (IFP) containing the Fas extracellular binding domain fused to a 4-1BB co-stimulatory domain, rather than the natural death domain. T cells engineered to express TCR_1045_ alone or in combination with the IFP were transferred into ID8_VEGF_-tumor bearing mice and evaluated for persistence, proliferation, anti-tumor cytokine production, and therapeutic efficacy.

**Results:** Relative to T cells modified only to express TCR_1045_, T cells engineered to express <u>both</u> TCR_1045_ and a Fas IFP preferentially persisted in the TME of tumor-bearing mice due to improved T cell proliferation and survival. Moreover, adoptive immunotherapy with IFP^+^ T cells significantly prolonged survival in tumor-bearing mice, relative to TCR_1045_ T cells lacking the IFP.

**Conclusions:** Fas/FasL signaling can mediate T cell death in the ovarian cancer microenvironment, as well as induce activation-induced cell death, an apoptotic mechanism responsible for regulating T cell expansion. Upregulation of FasL by tumor cells and tumor vasculature represents a mechanism for protecting growing tumors from attack by tumor-infiltrating lymphocytes. As many solid tumors overexpress FasL, an IFP that converts the Fas-mediated death signal into pro-survival and proliferative signals may provide an opportunity to enhance engineered adoptive T cell therapy against many malignancies.

## Introduction

More than 20,000 women are diagnosed with ovarian cancer annually, and >50% will die within 5 years^1^. This mortality rate has changed little in the last 20 years, highlighting the need for therapy innovation^2^. A rapidly evolving strategy with the potential to control tumor growth without toxicity to healthy tissues employs immune T cells engineered to selectively target proteins uniquely overexpressed in tumors. Mesothelin (MSLN) contributes to the malignant and invasive phenotype in ovarian cancer^3,4^, and has limited expression in healthy cells^5^, making it a candidate immunotherapy target in these tumors^6^. Studies targeting MSLN with monoclonal antibodies^7^, vaccination^8^, and chimeric antigen receptor (CAR)-expressing T cells^9^ have shown evidence of anti-tumor responses with acceptable safety profiles, validating MSLN as a viable target antigen.

In a murine ovarian cancer model of established, disseminated ID8_VEGF_ tumor, adoptively transferred T cells engineered to target Msln (TCR_1045_) preferentially accumulated within the established tumors, delayed tumor growth, and significantly prolonged survival^10^. However, this study also revealed that elements in the tumor microenvironment (TME) limit engineered T cell persistence and the ability to kill and eradicate cancer cells. Fas is expressed on antigen-experienced T cells, and FasL expression in the tumor vasculature of human and murine ovarian cancer can induce apoptosis of these Fas-expressing T cells^11,12^. To overcome this mechanism facilitating evasion from T cell responses, we generated a panel of immunomodulatory fusion proteins (IFP) containing the Fas receptor (Fas) extracellular binding domain fused to a 4-1BB co-stimulatory domain rather than the natural death domain^13^. Our prior results demonstrated this IFP can improve therapeutic efficacy of engineered T cells in mouse models of pancreatic cancer and acute myeloid leukemia^13^, but the mechanism of *in vivo* action and broader generalization of this strategy remained undefined.

We hypothesized that T cells expressing TCR_1045_ and a Fas-4-1BB IFP would have improved T cell persistence within ID8_VEGF_ tumors by disrupting apoptosis and enhancing T cell proliferation *in situ*. Our study reveals that the dominant mechanism by which the Fas-4-1BB IFP promotes T cell persistence *in vivo* is by increasing T cell survival within ovarian tumors. We also assessed the therapeutic activity of T cells engineered with TCR_1045_ and a Fas-4-1BB IFP and demonstrate improved survival of tumor-bearing mice without on-target, off-tumor toxicity. These studies validate that Fas-mediated apoptotic T cell death, an obstacle to the efficacy of engineered T cell therapy, can be addressed using a rationally designed fusion protein.

## Results

### FasL expression on ovarian tumor cells and effector T cells induces CD8 T cell death

FasL expression by solid tumors promotes tumor metastasis^14^ and immune evasion^15^. FasL is overexpressed in ovarian tumor vasculature^12^ and has higher expression in the ovarian TME than normal ovarian tissue^16^. We sought to evaluate the distribution of FasL expression in high grade serous ovarian cancer. Serial sections from matched primary and metastatic high grade serous ovarian cancer patient tumor samples were stained for CD31 and FasL (**Figure 1A**,**B**). FasL expression was detected above background in the vasculature of 12/17 primary and 14/17 metastatic tumors, consistent with previous reports^12,16^, but was also detected in the epithelial tumor regions of 11/17 primary and 13/17 metastatic tumors. Nearest neighbor Halo image analysis of the merged images revealed heterogeneous expression in the epithelial regions, with highest levels often found near the tumor edge (**Figure 1C**). Thus, T cells infiltrating ovarian tumors can encounter FasL death signals not only while exiting the vasculature but also throughout epithelial tumor regions.

**Figure 1.**
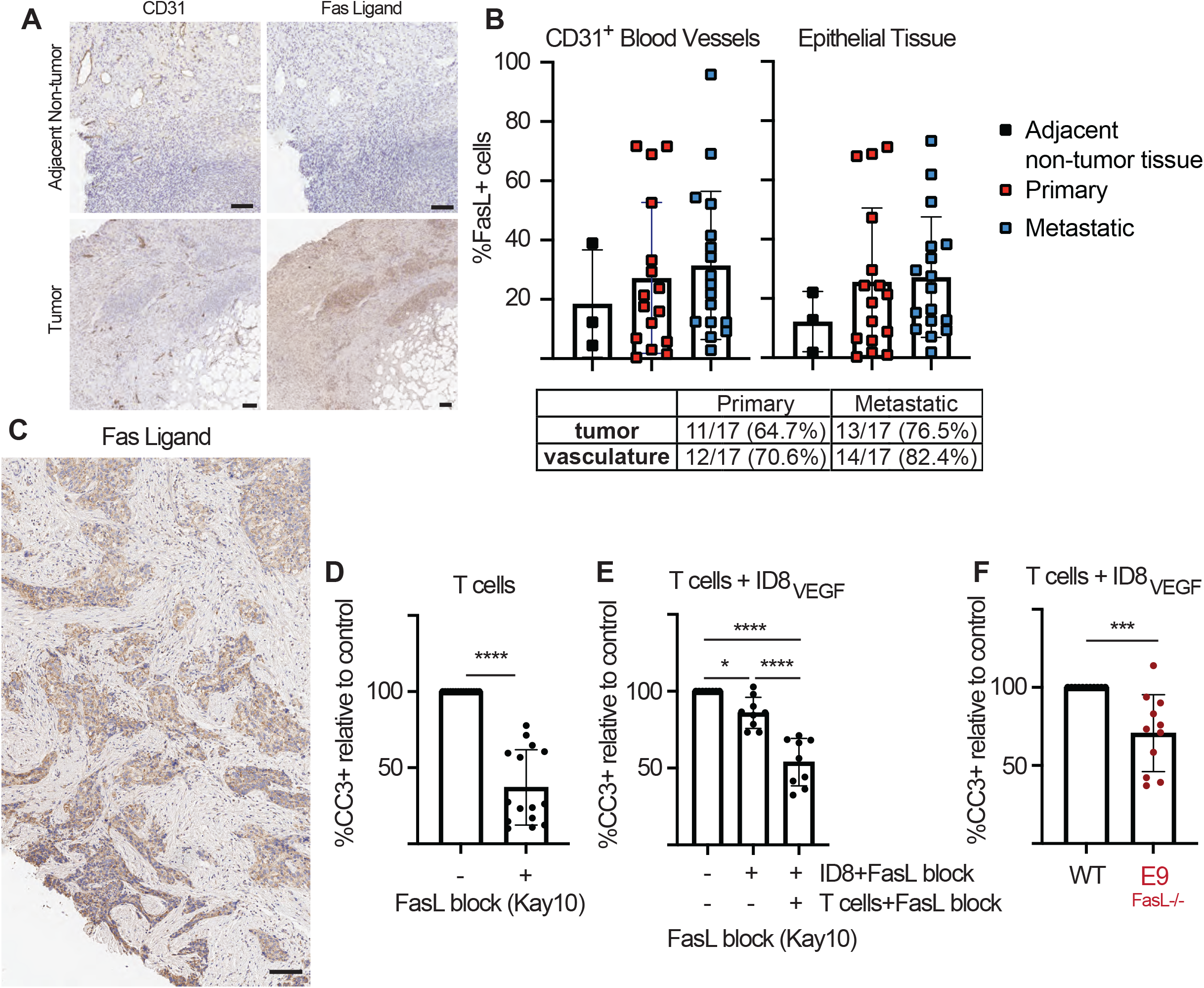
Fas Ligand signaling from T cells and ID8_VEGF_ tumor cells induce death in effector CD8 T cells. **A)** Immunohistochemistry staining for CD31 and Fas Ligand in human high grade serous ovarian cancer. Images are representative of 17 patient primary and 17 metastatic samples. Scale bar = 500 um. **B)** Halo quantification of FasL+ cells in CD31+ vasculature or tumor epithelium. **C)** Representative IHC staining for FasL, with higher intensity staining near the tumor edge. Scale bar = 100 um. **D)** Cleaved Caspase 3 (CC3) flow cytometry staining in activated CD8 T cells 3 days after culture with or without 24-hour pre-treatment with anti-FasL blocking antibody (clone: Kay10). Data are displayed as a percentage of CC3^+^ staining relative to untreated T cells. Cumulative data from 5 independent experiments. Student’s unpaired t-test. **** p<0.0001 **E)** CC3 flow cytometry staining in activated CD8 T cells 3 days after culture alone or co-culture with ID8_VEGF_ tumor cells. T cells or tumor cells were pre-treated with FasL (Kay10) blocking antibody for 24 hours as indicated. Data displayed as a percentage relative to T cells co-cultured with ID8_VEGF_ tumor cells without FasL blockade. Cumulative data from 3 independent experiments. One way ANOVA for multiple comparisons. * p=0.0353; **** p<0.0001 **F)** CC3 flow cytometry staining in activated CD8 T cells 3 days after co-culture with wild type ID8_VEGF_ or CRISPR edited ID8_VEGF_^FasL-/-^ tumor cells (E9 clone). Data displayed as a percentage relative to T cells co-cultured with wild type ID8_VEGF_ tumor cells. Cumulative data from 4 independent experiments. Student’s unpaired t-test. *** p=0.0008.

FasL is also expressed on activated T cells^17,18^. To evaluate if FasL signaling from fellow T cells can induce fratricide, we activated wild type P14 CD8 T cells, which express a TCR (TCR_gp33_) specific for the gp33-41 epitope of lymphocytic choriomeningitis virus (LCMV), *in vitro* with anti-CD3 and CD28 antibodies (Ab) for eight hours. We then treated the T cells with media or a FasL blocking Ab (Kay-10 clone) and stained the T cells for Cleaved Caspase 3 (CC3) three days later. FasL blockade significantly reduced CC3 expression (**Figure 1D**), suggesting T cells in culture engage in Fas/FasL signaling that induces apoptosis. To determine if FasL expression on ID8_VEGF_ tumor cells also induces CD8 T cell death, we co-cultured P14 CD8 T cells that had previously been activated *in vitro* with anti-CD3 and CD28 Ab, with wild type ID8_VEGF_ cells, with or without pre-treatment with FasL blocking Ab, at a 5:1 T cell to tumor cell ratio and stained for CC3 in T cells three days later. A significant decrease in CC3 staining was observed when FasL signaling from tumor cells was inhibited by the Ab (**Figure 1E**), which was further reduced when FasL from T cells was blocked concurrently. To verify the results of the blocking assay, we used CRISPR/Cas9 targeting to generate a FasL knockout ID8_VEGF_ cell line (ID8_VEGF_^FasL-/-^; denoted E9; **Supplemental Figure 1**). T cells activated with anti-CD3 and CD28 Ab were co-cultured with wild type ID8_VEGF_ or E9 ID8_VEGF_^FasL-/-^ cells, and CC3 staining was reduced in the T cells that were co-cultured with ID8_VEGF_^FasL-/-^ cells (**Figure 1F**), confirming that effector CD8 T cells are susceptible to FasL signals from both ID8_VEGF_ tumor cells and fellow CD8 T cells.

### TCR_1045_/Fas-4-1BB IFP^+^ T cells exhibit enhanced survival *in vivo*

We previously isolated a high-affinity murine TCR specific for the Msln_406-414_ epitope (designated TCR_1045_) and demonstrated CD8 T cells transduced with TCR_1045_ mediate anti-tumor activity against ovarian and pancreatic cancer without toxicity to normal tissues^10,19^. We also previously developed an immunomodulatory fusion protein (IFP) containing the Fas receptor (Fas) extracellular binding domain fused to a 4-1BB co-stimulatory domain and showed that expression of this fusion protein improved the efficacy of T cell therapy in mouse models of acute myeloid leukemia and pancreatic cancer^13^. We now sought to evaluate if the Fas-4-1BB_tm_ IFP could improve therapeutic activity in a mouse model of ovarian cancer in which FasL/Fas signaling is known to be an obstacle to efficacy^12^. Naive transgenic P14 CD8 T cells were activated with anti-CD3 and CD28 Ab and transduced with either a bi-cistronic vector encoding the alpha and beta chains of TCR_1045_ or a tri-cistronic vector encoding both TCR_1045_ chains and a Fas-4-1BB_tm_ IFP (TCR_1045_/Fas-4-1BB_tm_, **Figure 2A**), linked by P2A elements to ensure equimolar expression. P14 T cells engineered with either TCR_1045_ or TCR_1045_/Fas-4-1BB_tm_ express similar levels of the Vβ9 component of TCR_1045_, but only TCR_1045_/Fas-4-1BB_tm_ T cells expressed high levels of Fas (CD95), reflecting high levels of the transduced IFP (**Figure 2B**). We previously reported that T cells expressing the Fas-4-1BB_tm_ construct produce increased levels of IL-2^13^, and we validated that the construct including both TCR_1045_ and the Fas-4-1BB_tm_ fusion protein also resulted in increased IL-2 production by transduced T cells (**Figure 2C**).

**Figure 2.**
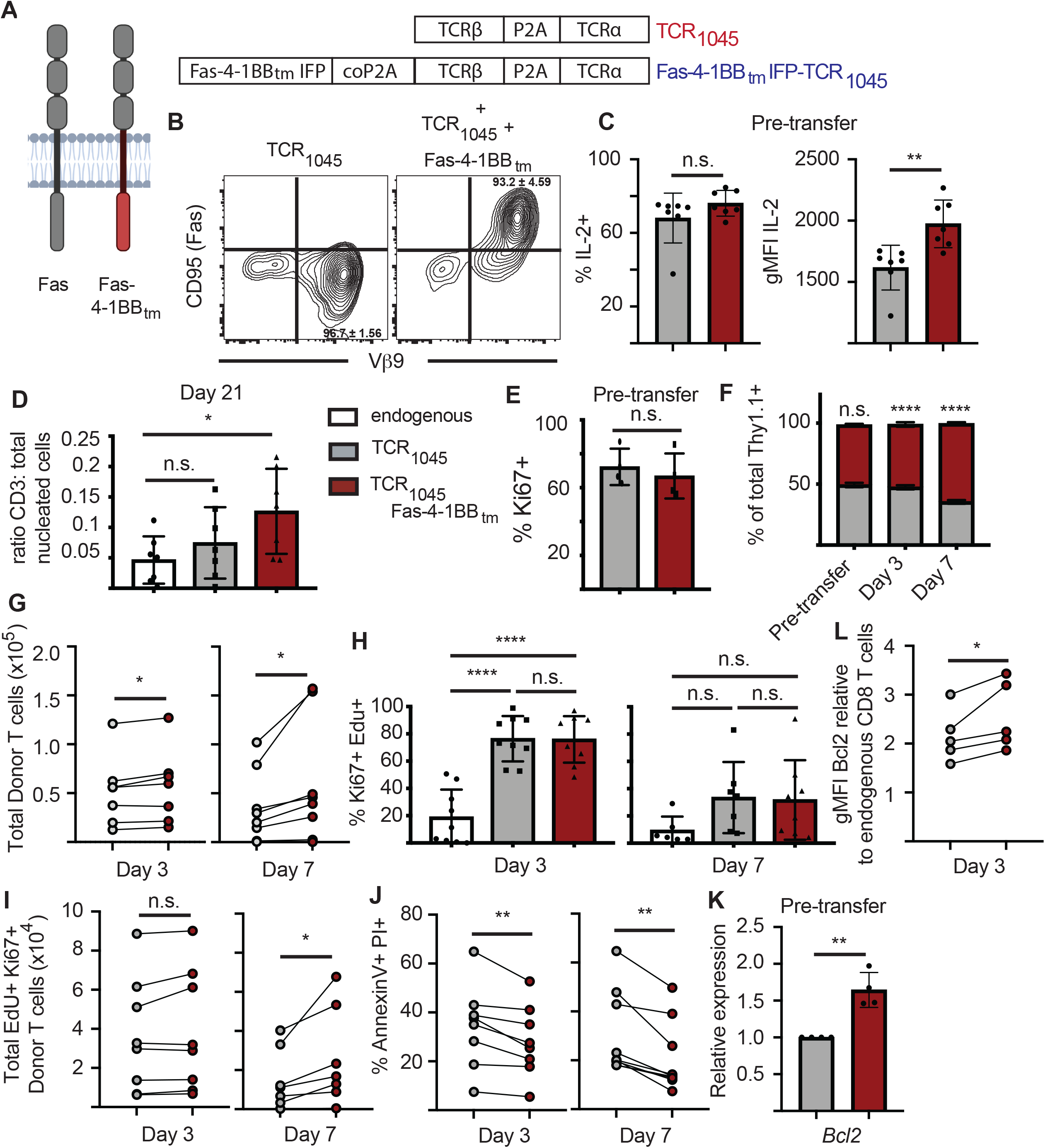
Expression of a Fas-IFP enhances accumulation of CD8 T cells *in vivo*. **A)** Schematic of monomeric endogenous Fas receptor and Fas-4-1BB immunomodulatory fusion proteins and vectors encoding TCR_1045_ or TCR_1045_ and a Fas-4-1BB_tm_ immunomodulatory fusion protein (IFP). Genes are separated by P2A or codon-optimized (co) P2A elements to ensure equimolar expression. **B)** CD95 (Fas) and Vβ9 expression on transduced P14 Thy1.1^+^ CD8 T cells. T cells were transduced with retroviral vectors containing TCR_1045_ or TCR_1045_ and Fas-4-1BB_tm_ IFP and re-stimulated with irradiated splenocytes pulsed with Msln_406-414_ in the presence of IL-2. Cells were screened by flow cytometry 7 days after restimulation. Data representative of >5 independent experiments. Quadrant values indicate mean and standard deviation from 5 independent experiments. **C)** Intracellular cytokine staining for IL-2 production by P14 Thy1.1^+^ CD8 T cells expressing either TCR_1045_ (gray) or TCR_1045_ and Fas-4-1BB_tm_ IFP (red) after 5 hour stimulation with Msln_406- 414_ peptide. Cumulative data from 3 independent experiments (1-3 technical replicates per experiment). Unpaired t-test, ** p=0.0040. n.s. = not significant. Error bars indicate standard deviation. **D)** IHC staining for CD3^+^ T cells in ID8_VEGF_ tumors 21 days after T cell transfer. P14 Thy1.1^+^ CD8 T cells expressing TCR_1045_ or TCR_1045_ and Fas-4-1BB_tm_ IFP were injected i.p. into ID8_VEGF_ tumor-bearing mice. Mice were pre-treated with 180mg/kg cyclophosphamide >6 hours prior to cell transfer. All mice received 1×10^7^ engineered T cells, 5×10^7^ irradiated splenocytes pulsed with Msln_406-414_ peptide, and daily IL-2 s.c. for 10 days (1×10^4^ IU). CD3 staining in untreated mice (white bar) included as a reference for endogenous T cell infiltration. Cumulative data from 3 independent experiments, n=7 per group. One-way ANOVA Tukey’s multiple comparisons test, * p=0.0442. n.s. = not significant **E)** Intracellular flow cytometry staining for Ki67 expression in Thy1.1^+^ CD8^+^ Vβ9^+^ T cells 7 days after *in vitro* restimulation of engineered T cells. Data from 4 independent experiments. Unpaired t-test. **F - L)** Thy1.1^+^ or Thy1.1/1.2^+^ P14 T cells expressing TCR_1045_ (gray) or TCR_1045_ and Fas-4-1BB_tm_ IFP (red) were co-transferred at a 1:1 ratio into ID8_VEGF_ tumor-bearing mice with 5×10^7^ irradiated splenocytes pulsed with Msln_406-414_ peptide, and daily s.c. IL-2. At 3 or 7 days after co-transfer, T cells were isolated from tumors, enumerated and evaluated by flow cytometry for indicated markers. **F)** Proportion of donor T cells within ID8_VEGF_ tumors after co-transfer. Data pooled from 3 independent experiments, n=8 (day 3) or n=7 (day 7) mice per group. 2-way ANOVA for multiple comparisons. **** p<0.0001. **G)** Quantification of intratumoral donor T cells. Counts of the co-transferred T cell populations from each individual mouse are connected with a line. Paired t-test. * p<0.05. **H)** Flow cytometry staining of intratumoral endogenous and donor T cells 3 or 7 days after transfer for Ki67 expression and EdU incorporation. EdU was injected i.p. into mice 24 hours prior to T cell isolation. 1-way ANOVA for multiple comparisons. **** p<0.0001. n.s. = not significant. **I)** Quantification of EdU+ Ki67+ intratumoral donor T cells. Counts of the co-transferred cell populations from each individual mouse are connected with a line. Paired t-test. * p<0.05. **J)** Flow cytometry staining of intratumoral donor T cells 3 or 7 days after transfer for Annexin V and Propidium Iodide (PI). Cell populations from the same mouse are connected with a line. Data from three independent experiments, n=8 (day 3) or n=9 (day 7). Paired t-test. ** p<0.01. **K)** qPCR for *Bcl2* expression in TCR_1045_ (gray) or TCR_1045_/Fas-4-1BB_tm_ IFP (red) T cells 8 days after restimulation *in vitro*. Data pooled from 4 independent experiments. Unpaired t-test ** p=0.0016. **L)** Flow cytometry staining of intratumoral engineered T cells 3 days after transfer for intracellular Bcl2. Cell populations from the same mouse are connected with a line. Data from three independent experiments, n=5. Paired t-test. * p=0.0376. All error bars represent SD.

We previously demonstrated that, although TCR_1045_ T cells infiltrate ID8_VEGF_ tumors, the antitumor activity is limited by failure of the T cells to persist^10^. To evaluate if TCR_1045_/Fas-4-1BB_tm_ T cells would persist more effectively in mouse ovarian tumors than control TCR_1045_ cells, T cells were transferred into ID8_VEGF_ tumor-bearing mice following a previously established protocol^10^. Briefly, 5×10^6^ ID8_VEGF_ cells were injected intraperitoneally (i.p.) into eight-week-old female C57Bl/6 mice. After tumor nodules were detected by high resolution ultrasound (>6 weeks after tumor injection), mice were treated with cyclophosphamide ≥6 hours prior to T cell transfer, and then received 1×10^7^ engineered T cells and a vaccine of 5×10^7^ irradiated splenocytes pulsed with Msln_404-416_ peptide to enhance engraftment^10^. Prior to transfer, >93% of the transduced T cells expressed TCR_1045_ or TCR_1045_/Fas-4-1BB_tm_, as indicated by staining for the Vβ9 component of the Msln-specific TCR (**Figure 2B**). By twenty-one days after T cell transfer, the number of TCR_1045_ T cells within tumors was not greater than the number of endogenous T cells in tumors from untreated mice, but TCR_1045_/Fas-4-1BB_tm_ T cells remained at 3-fold greater numbers than T cells in untreated tumors (**Figure 2D**). Co-stimulation through 4-1BB promotes T cell proliferation and survival^20,21^, thus the enhanced persistence of TCR_1045_/Fas-4-1BB_tm_ T cells within tumors could be due to increased T cell proliferation, reduced T cell death or both. To evaluate if T cell proliferation was enhanced *in vivo*, we evaluated Ki67 expression and EdU incorporation by tumor-infiltrating engineered T cells. Congenically distinct (Thy1.1^+^ or Thy1.1^+^Thy1.2^+^) P14 T cells were transduced with TCR_1045_ or TCR_1045_/Fas-4-1BB_tm_ and equal numbers of transduced cells were co-transferred into tumor-bearing mice. Prior to T cell transfer, TCR_1045_ and TCR_1045_/Fas-4-1BB_tm_ T cells expressed similar levels of Ki67 (**Figure 2E**), suggesting that expression of the Fas-4-1BB_tm_ construct does not change the proportion of activated cells in cell cycle. At three or seven days after transfer, TCR_1045_/Fas-4-1BB_tm_ preferentially accumulated within tumors (**Figure 2F**,**G**). Although there was no difference in the proportion of Ki67^+^Edu^+^ cells at either time point (**Figure 2H**), we observed 1.7-fold more Ki67^+^EdU^+^ TCR_1045_/Fas-4-1BB_tm_ T cells seven days after transfer (**Figure 2I**). While this may suggest enhanced proliferation *in situ*, this difference could also be explained by reduced cell death in the TCR_1045_/Fas-4-1BB_tm_ T cell population. To determine if TCR_1045_/Fas-4-1BB_tm_ T cells do exhibit enhanced survival, we evaluated Annexin V and Propidium Iodide (PI) staining of intratumoral T cells. Three or seven days after transfer, 1.2-fold and 1.5-fold fewer TCR_1045_/Fas-4-1BB_tm_ T cells than TCR_1045_ T cells, respectively, were Annexin V^+^/PI^+^ (**Figure 2J**), suggesting the dominant mechanism of improved TCR_1045_/Fas-4-1BB_tm_ T cell persistence is enhanced survival within ID8_VEGF_ tumors.

To further interrogate the mechanism of TCR_1045_/Fas-4-1BB_tm_ T cell persistence, expression of the pro-survival protein Bcl2 was quantified by qPCR eight days after activation *in vitro* (**Figure 2K**) and was detected at higher levels in TCR_1045_/Fas-4-1BB_tm_ T cells relative to TCR_1045_ T cells. Intracellular staining of TCR_1045_/Fas-4-1BB_tm_ T cells isolated from tumors three days after transfer also revealed higher Bcl2 expression than in TCR_1045_ T cells (**Figure 2L**). Together, these data suggest that TCR_1045_/Fas-4-1BB_tm_ T cells achieve enhanced persistence in ID8_VEGF_ tumors predominantly by improved T cell survival within the TME.

### TCR_1045_/Fas-4-1BB IFP^+^ T cells sustain enhanced IL2 production in solid tumors

We previously reported that expression of the Fas-4-1BB_tm_ IFP enhanced cytokine production *in vitro*^13^. To evaluate if this advantage would be maintained *in vivo*, we co-transferred P14 T cells expressing TCR_1045_ or TCR_1045_/Fas-4-1BB_tm_ into tumor-bearing mice, as previously described. Twenty-one days after cell transfer, both tumor-infiltrating T cell populations expressed PD-1, Lag-3, Tim-3, and TIGIT (**Figure 3A**, relative to naïve T cells in the spleen). Fas-4-1BB_tm_ IFP^+^ T cells expressed reduced PD-1 and elevated Tim-3 and Lag-3 relative to their TCR_1045_ counterparts. Consistent with our previous findings, engineered T cells isolated from tumors and stimulated with Msln_406-414_ peptide *ex vivo* produced less of the effector cytokines interferon-γ (IFNγ) and tumor necrosis factor-α (TNFα) than cells isolated from the spleen^10^, but no differences were detected between TCR_1045_ and TCR_1045_/Fas-4-1BB_tm_ T cells (**Figure 3B**). However, consistent with our *in vitro* results (**Figure 2C**), TCR_1045_/Fas-4-1BB_tm_ T cells from the tumor produced more IL-2 than TCR_1045_ T cells (**Figure 3C**). This further suggests that FasL signals from the TME may be promoting sustained 4-1BB co-stimulation through the IFP, supporting T cell survival and proliferation. These data also demonstrate that TCR_1045_ and TCR_1045_/Fas-4-1BB_tm_ T cells are both potentially susceptible to inhibitory signals present in the TME.

**Figure 3.**
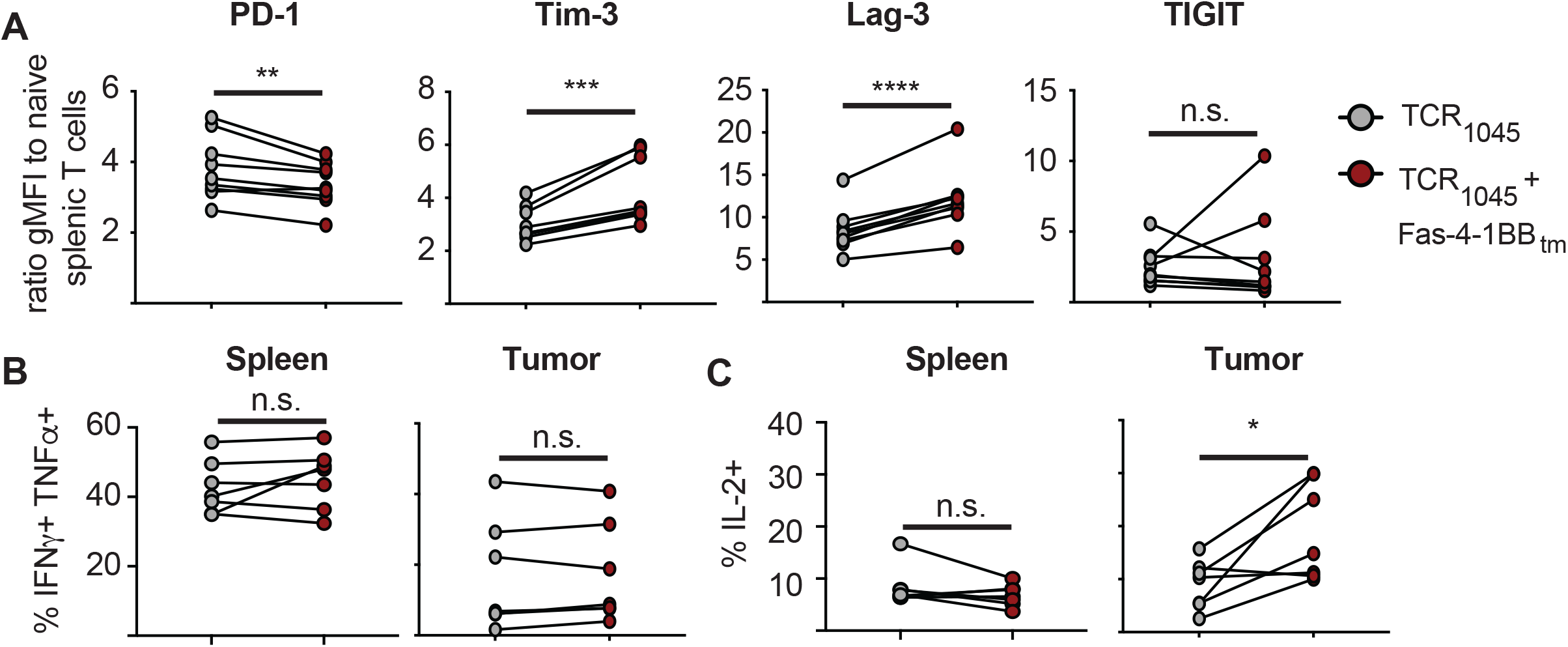
Fas-IFP-expressing T cells remain susceptible to T cell dysfunction *in vivo*. Thy1.1^+^ or Thy1.1/1.2^+^ P14 T cells expressing TCR_1045_ or TCR_1045_/Fas-4-1BB_tm_ IFP were co-transferred at a 1:1 ratio i.p. into ID8_VEGF_ tumor-bearing mice with 5×10^7^ irradiated splenocytes pulsed with Msln_406-414_ peptide, and daily s.c. IL-2. **A)** At 21 days after co-transfer, T cells were isolated from tumors and evaluated by flow cytometry for expression of PD-1, Tim-3, Lag-3, and TIGIT. TCR_1045_ T cells (gray) and TCR_1045_/Fas-4-1BB_tm_ IFP^+^ T cells (red) from the same mouse are connected with a line. Data are displayed as a ratio of the geometric mean fluorescence intensity (gMFI) of Thy1.1^+^ T cells compared to endogenous CD44^-^ CD62L^+^ naïve CD8 T cells from the spleen of the same mouse. Cumulative data from 2 independent experiments (n=7 mice). Paired t-test. ** p=0.0064; *** p=0.0005; **** p<0.0001; n.s.=not significant. **B and C)** *Ex vivo* intracellular cytokine staining for IFN-γ and TNF-α or IL-2 production by P14 Thy1.1^+^ TCR_1045_ (gray) or TCR_1045_/Fas-4-1BB_tm_ IFP (red) CD8 T cells isolated from spleen or tumor, after 5-hour stimulation with Msln_406-414_ peptide. Cumulative data from 2 independent experiments (n=7 mice). Paired t-test. * p<0.05

### Adoptive immunotherapy with TCR_1045_/Fas-4-1BB_tm_ T cells significantly prolongs survival of ovarian tumor-bearing mice

To evaluate if TCR_1045_/Fas-4-1BB_tm_ T cells control growth of ovarian cancer more effectively than TCR_1045_ T cells, we used an established treatment protocol that previously demonstrated TCR_1045_ T cells prolong survival of ID8_VEGF_ tumor-bearing mice^10^. Briefly, ID8_VEGF_ tumor-bearing mice were treated with a single dose of cyclophosphamide ≥6 hours prior to T cell transfer, and then received 1×10^7^ engineered T cells and 5×10^7^ irradiated splenocytes pulsed with Msln_404-416_ peptide every 14 days. Mice received 10^4^ U IL-2 daily for 10 days after each T cell transfer to promote T cell expansion and persistence. Mice treated with TCR_1045_/Fas-4-1BB_tm_ T cells had a significantly longer median survival (123 days) relative to control mice (77 days) or mice treated with TCR_1045_ T cells (102 days) (**Figure 4A**).

**Figure 4.**
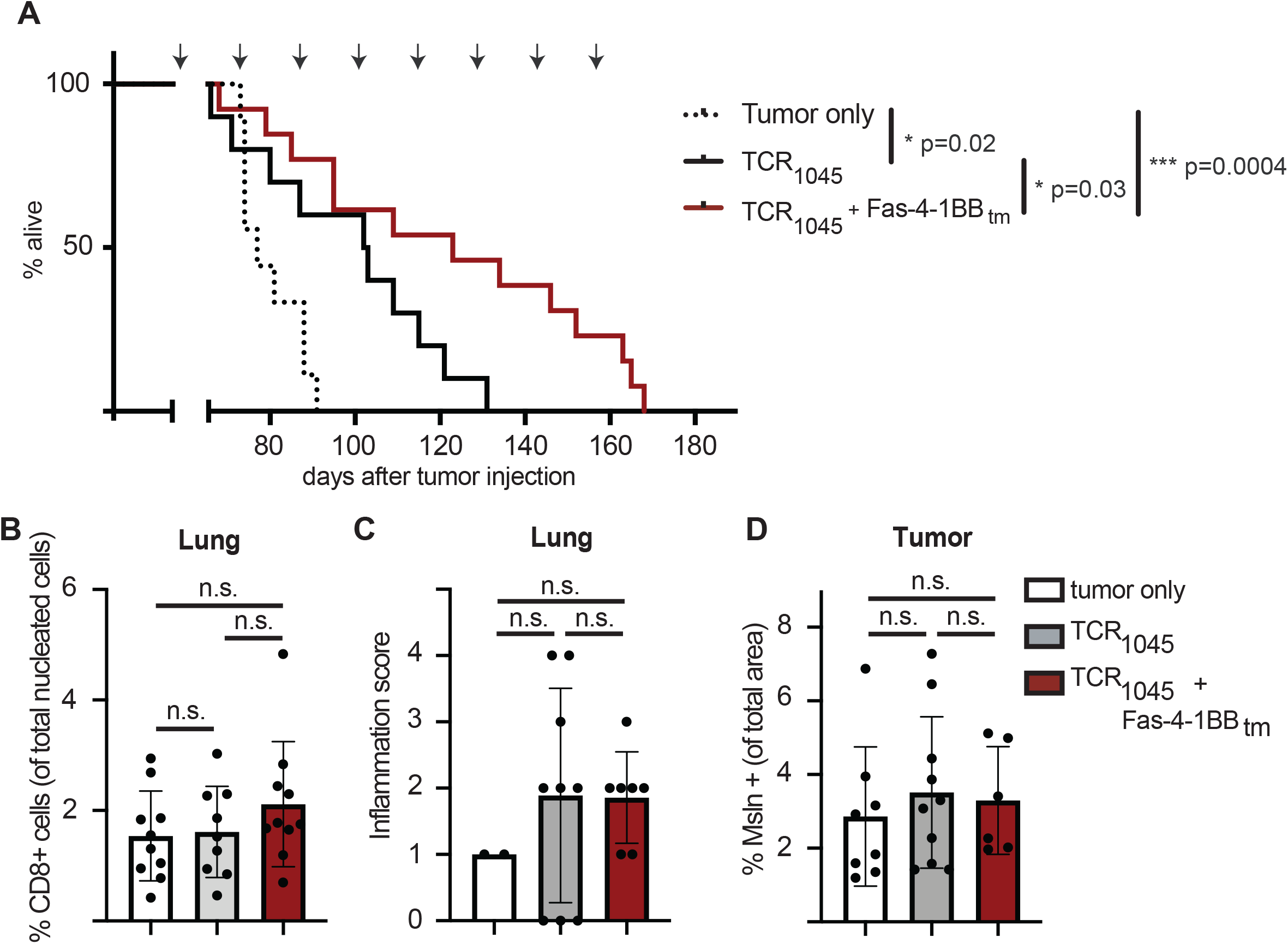
A Fas-IFP enhances anti-tumor efficacy *in vivo*. **A)** Overall survival of ID8_VEGF_ tumor-bearing mice treated with repeated doses of P14 Thy1.1^+^ CD8 T cells transduced with TCR_1045_ or TCR_1045_ and Fas-4-1BB_tm_ IFP. Tumor-bearing mice received a single dose of cyclophosphamide (180mg/kg) before the 1^st^ T cell infusion followed by injections of 1×10^7^ TCR_1045_ or TCR_1045_/Fas-4-1BB_tm_ T cells and peptide-pulsed irradiated splenocytes (5:1 APC:T cell ratio) i.p. every 14 days with 1×10^4^ U IL-2 s.c. for 10 days after each infusion. Treatment was initiated 45-52 days after tumor injection, when tumors were detectable by US. Arrows above survival curve indicate the timing of T cell infusions. Survival data are aggregated from 3 independent experiments, n=9-13 total per group. Log-rank (Mantel-Cox) test. **B)** IHC quantification of CD8+ T cells in lungs of untreated (white), TCR_1045_ (gray) or TCR_1045_/Fas-4-1BB_tm_ IFP (red) treated mice at necropsy. **C)** Histology scoring of hematoxylin and eosin IHC stained lungs from untreated (white), TCR_1045_ (gray) or TCR_1045_/Fas-4-1BB_tm_ IFP (red) treated mice at necropsy. All lungs contained metastatic tumor. Scoring ranged from 0 (tissue within normal limits), to 4 (neutrophilic interstitial pneumonia with or without lymphoplasmacytic and neutrophilic perivasculitis) and was performed by a trained pathologist. **D)** IHC quantification of Msln in tumors of untreated (white), TCR_1045_ (gray) or TCR_1045_/Fas-4-1BB_tm_ IFP (red) treated mice at necropsy. B-D) Cumulative data from 3 independent experiments, n=9-10 per group, except the untreated mice in C), for which only 2 mice had metastatic tumors in the lungs. One-way ANOVA with multiple comparisons. All error bars represent SD.

Since Msln is expressed on the lining of the pleural and pericardial cavities, we evaluated if expression of the IFP resulted in enhanced T cell accumulation in the lungs. At euthanasia, the lungs of treated and untreated mice were evaluated for CD8 T cell infiltration by IHC staining and the proportions of CD8 T cells present in the lungs from the treatment groups were not statistically different (**Figure 4B**).

To assess inflammation, H&E-stained sections of the lungs and hearts of treated and untreated mice were evaluated for metastatic tumor growth and neutrophil infiltration by a pathologist. No treatment-related myocarditis or cardiac lesions were observed. We detected metastatic tumor growth in lung samples from 2/7 untreated mice, 9/9 TCR_1045_ T cell-treated mice, and 7/12 TCR_1045_/Fas-4-1BB_tm_ T cell-treated mice, with lungs from mice that survived the longest harboring the largest metastatic nodules (Note: samples with poor fixation or staining were excluded from the analysis). No inflammation was observed in the lungs of mice without metastatic lesions. Mice with metastatic tumor growth in the lungs were scored to determine if the presence of metastases in the lung increased inflammation after T cell treatment. Mild to moderate inflammation, defined as lymphoplasmacytic and neutrophilic perivasculitis, was observed in the lungs of some treated mice (**Figure 4C**) and was scored slightly higher in treated compared to untreated mice, but this did not reach statistical significance. No mice showed signs of respiratory distress during the course of the experiment.

To determine if mice that received TCR_1045_/Fas-4-1BB_tm_ T cells succumbed to disease due to loss of expression of the target antigen, the tumors of treated and untreated mice were evaluated for Msln expression by IHC. No differences were observed (**Figure 4D**), suggesting that antigen loss has not occurred in the treated tumors, and consistent with the upregulation of exhaustion markers we observed on tumor-infiltrating cells (**Figure 3A**).

## Discussion

Fas/FasL signaling can mediate T cell death, including activation-induced cell death, an apoptotic mechanism responsible for regulating T cell expansion during repeated stimulation, suggesting tumor cells may upregulate FasL for protection from tumor-infiltrating lymphocytes. Here we have shown that relative to T cells modified with only TCR_1045_, T cells engineered to express <u>both</u> TCR_1045_ and a Fas-4-1BB IFP preferentially persist in the ovarian TME following transfer into tumor-bearing mice. Moreover, adoptive immunotherapy with IFP^+^ T cells significantly prolonged survival in tumor-bearing mice, relative to TCR_1045_ T cells lacking an IFP. Our findings suggest that the dominant mechanism supporting the improved T cell persistence within tumors is enhanced T cell survival; however, since the IFP^+^ T cells produce more IL-2 than their IFP^-^ counterparts and a greater number of EdU^+^ IFP^+^ cells were detected, the IFP^+^ T cells may also undergo greater proliferation *in vivo* that was not captured in the two timepoints we evaluated using EdU incorporation.

The Fas-4-1BB IFP utilizes a decoy Fas receptor to provide co-stimulatory signals, which are often limited in solid tumors. Cell-intrinsic engineering strategies that truncate the Fas receptor have been shown to successfully disrupt death receptor signaling and improve the efficacy of T cell therapy in a mouse model of melanoma^16^, but such approaches providing a dominant negative Fas molecule still lack co-stimulatory signals that may be critical for T cell persistence in some TME. Systemic agents that provide co-stimulation, such as 4-1BB agonists and soluble 4-1BB ligand, have been shown to promote anti-tumor immunity, although these agents are not cell intrinsic and can also trigger high levels of inflammation^22^. The Fas-4-1BB IFP targets co-stimulation specifically to tumor-specific T cells, resulting in intratumoral T cells that have greater expression of anti-apoptotic molecules and production of IL-2, which in concert promote T cell survival and enhance the efficacy of T cell therapy. As FasL is expressed at high levels in many solid tumors,^16^ which is associated with poor prognosis^11^, the use of Fas-4-1BB IFPs may provide an opportunity to enhance engineered adoptive T cell therapy for many malignancies.

## Methods

### Human specimens

All studies using human specimens were approved by the Fred Hutchinson Cancer Research Center Institutional Review Board and conducted according to the principles expressed in the Declaration of Helsinki. Tumor tissues were obtained by the POCRC Repository from patients who provided written informed consent.

### Cell Lines

ID8_VEGF_ cells, which were transduced to overexpress vascular endothelial growth factor (VEGF) to recapitulate the elevated levels observed in human disease^23^, were a gift from Dr. Matthias Stephan in 2014. ID8 cells were cultured in DMEM (Gibco) containing 10% FBS (Hyclone), 1% Penn/Strep (Gibco), and 0.1% Insulin-Transferrin-Selenium (Sigma). ID8_VEGF_ cell lines were passaged fewer than 10 times before use in experiments. PLAT-E cells (ATCC) passaged fewer than 35 times were used for retrovirus production for all T cell transductions. All cell lines were confirmed negative for mycoplasma prior to use. Cell lines were not authenticated in the past year.

### Mouse strains

The Institutional Animal Care and Use Committees of the University of Washington and the Fred Hutchinson Cancer Research Center approved all animal studies. 5×10^6^ ID8_VEGF_^23,24^ cells were injected i.p. into 6-7 week old female C57BL/6J mice (Jackson Laboratories). P14 mice have been previously described^25^.

### ID8_VEGF_ FasL CRISPR KO

ID8_VEGF_ cells were edited using CRISPR knockout technology by Synthego. Briefly, founder ID8_VEGF_ cells (mycoplasma-free) were sequenced and synthetic single guide RNA (sgRNA) was designed to target FasL (guide target sequence: CTCCTTTGGTCCGGCCCTCT). Specific guide RNA was complexed with *Streptococcus pyogenes* Cas9 (PAM motif: AGG) into a ribonucleoprotein and delivered into ID8_VEGF_ cells by electroporation. PCR-amplification and Sanger sequencing was used to sequence the transfected cells to verify knockout efficiency. Cells from the edited pool were seeded using single cell dilution and expanded to establish clonal lines. Clones were sequenced to verify the clone contained a homozygous edit and that the progeny were derived from a single cell, and a single clone (E9) was selected for experiments.

### Cytotoxicity Assays

Naïve P14 T cells were activated for 72 hours with anti-CD3 and CD28 antibodies, then cultured alone or co-cultured with tumor cells for 18-24 hours in 24-well tissue culture plates. T cells were then gently removed and transferred to FACS tubes for intracellular cleaved-caspase 3 (CC3) staining. The BD Fix/Perm kit (cat: 554714) was used for staining with anti-CC3.

### Retroviral Constructs

The codon-optimized murine Vβ9 and Vα4 TCR chains, connected by a porcine teschovirus-1 2A element (“P2A”; Life technologies), recognize the Msln_406-414_ epitope presented in H2-D^b^, and were cloned from the Mig-R1 retroviral vector^19^ into the pENTR vector and subsequently Gateway cloned into the pMP71 retroviral vector for all TCR_1045_ studies. A construct encoding Fas-4-1BB_tm_ and TCR_1045_ was ordered from GeneArt (ThermoFisher) in the pDONR221 vector and gateway cloned into the pMP71 retroviral vector for all TCR_1045_ studies.

### TCR transduction of T cells

T cell transduction has been described previously^10^ and can be accessed at DOI: <u>dx.doi.org/10.17504/protocols.io.smrec56</u>. Briefly, PLAT-E cells were transfected with DNA encoding TCR_1045_ or TCR_1045_/Fas-4-1BB_tm_, the media was replaced at 24 hours, and the viral supernatant used for T cell transduction 72 and 96 hours. Splenic T cells isolated from P14 Thy1.1^+^ mice were stimulated with anti-CD3 and anti-CD28 antibodies in the presence of IL-2 and transduced with retroviral supernatant by spinfection in polybrene at 24 and 48 hours after activation. T cell restimulation has been described previously^10^ and can be access at DOI: <u>dx.doi.org/10.17504/protocols.io.spqedmw</u>. Briefly, Thy1.2^+^ splenocytes were irradiated and pulsed with Msln_406-414_ peptide for >90 minutes to prepare APCs. Transduced T cells were co-cultured with peptide-pulsed APCs for 5-7 days in the presence of IL-2. T cells were screened by flow cytometry for transduction efficiency 5 days after activation or restimulation.

### T cell isolation

Tumors were dissociated in 3ml of RPMI with 10% FBS using the m_imptumor_01 setting on a gentleMACS dissociator (Miltenyi Biotec). Samples were transferred to a conical tube with 35mL of Collagenase (RPMI containing 1% HG solution, 0.1% MgCl2, 1%CaCl2, 5% FBS, and 20,000U of Collagenase Type IV) and mixed on a MACSmix rotator at 37°C for 10 minutes. Samples were filtered through cell strainers (70μm Falcon) and lymphocytes were purified on a 44/67% percoll gradient (800x*g* at 20°C for 20 minutes). Spleens were dissociated by mechanical separation through a cell strainer and ACK lysis (Gibco, cat: A10492-01) was performed to remove red blood cells.

### Flow cytometry

A MSLN_406-414_/H2-D^b^ tetramer conjugated to APC was prepared by the Fred Hutch Immune Monitoring Core. All cells were stained with LIVE/DEAD fixable Aqua (405nm, cat: L34966) in 1x DPBS prior to surface or intracellular staining. UltraComp eBeads (eBioscience, cat: 01-2222) were used for all compensation. For *ex vivo* experiments, cells from either untreated mice or endogenous CD44^-^ CD62L^+^ CD8^+^ T cells from the spleen of treated mice were used for negative controls and gating. For *in vitro* experiments, fluorescence minus one or irrelevant engineered T cells were used for negative controls and gating. The eBioscience Fix/Perm kit (cat: 00-5523-00) was used for staining with anti-Ki67 (cat: 51-36524X) with a mouse IgG1 isotype as a control (cat: 51-35404X). Cells were resuspended in FACS buffer (1x DPBS containing 2% FBS Hyclone and 0.72% 0.5M EDTA) or 0.5% Paraformaldehyde and acquired with an LSR2-2 (BD).

### Intracellular cytokine stimulation

The stimulation protocol for intracellular cytokine staining has been described previously^10^ and can be found at DOI: <u>dx.doi.org/10.17504/protocols.io.sqdeds6</u>. Briefly, cells were treated with protein transport inhibitor containing Brefeldin A (GolgiPlug) and plated at 1×10^6^ cells per well in a flat-bottom 96-well plate. Cells were stimulated for 5 hours at 37C with T cell media or 1 mg/ml of Msln_406-414_ peptide (GQKMNAQAI). Peptides were ordered from ELIM peptide (>80% purity). The BD Fix/Perm kid was used for intracellular staining. In some experiments, cells were fixed in 0.5% paraformaldehyde until data acquisition.

### Cell proliferation

The Click-iT Plus EdU imaging kit (cat: C10640) was reconstituted at 10mM in DMSO and used to analyze *in situ* cell proliferation. For *in vitro* experiments, 3×10^6^ cells were plated with 5uL of the 10mM EdU solution and incubated for 24 hours. Cells were treated with the eBioscience Fix/Perm kit and subsequently stained using Click-iT Plus EdU imaging kit reagents: Click-iT reaction buffer, copper protectant, Alexa Fluor picolyl azide (AF647 fluorescent), and Click-iT reaction buffer additive. Samples were analyzed by flow cytometry within 4 hours. For *in vivo* EdU experiments, EdU was diluted to 5mg/mL of saline. Mice were injected i.p. with 50mg EdU/kg and cells were isolated 24 hours later by enzymatic digestion. Cells were stained using the Click-iT Plus EdU imaging kit and analyzed by flow cytometry within 4 hours.

### Quantitative PCR

3×10^6^ transduced T cells were resuspended in RLT lysis buffer (supplemented with b-ME) and RNA extracted with the Qiagen RNeasy Plus RNA isolation kit. RNA integrity was analyzed on Tape Station analyzer and cDNA was generated using iScript Reverse Transcription Supermix for RT-qPCR (Biorad). qPCR was run using qPCR SYBR Green Assay with mouse BCL2 PrimePCR assay (Biorad) and with RPL13a reference gene (Primers: Fwd.: TTCTCCTCCAGAGTGGCTGT, Rev.: GGCTGAAGCCTACCAGAAAG) on the 384 well ABI QuantStudio5 instrument. The delta delta CT method was used for analysis.

### Adoptive immunotherapy

ID8_VEGF_-tumor-bearing mice received either engineered T cells (1×10^7^, transduced and re-stimulated *in vitro*) or engineered T cells (1×10^7^) transduced without restimulation *in vitro* but then stimulated *in vivo* with 5×10^7^ peptide-pulsed irradiated splenocytes as a vaccine. Cell infusions were followed by IL-2 (2×10^4^ IU, s.c.) daily for 10 days to promote T-cell expansion and survival. For therapy with serial T-cell infusions, mice received this same treatment protocol every two weeks. In indicated experiments, treated mice received one dose of Cyclophosphamide (180 mg/kg) i.p. to lymphodeplete hosts approximately 6-8 hours prior to only the first T-cell transfer.

### Immunohistochemistry

Histology preparation has been described previously^10^ and can be found at DOI: <u>dx.doi.org/10.17504/protocols.io.sppedmn</u>. Briefly, harvested tissues were fixed in 10% neutral buffered formalin for at least 72 hours, embedded in paraffin, sectioned (4um) and stained with hematoxylin and eosin or primary antibodies for markers of interest using a Leica Bond Instrument. Following antigen retrieval, slides were blocked with Leica Bond Peroxide Block and then 10% normal goat serum, stained with primary antibodies for 30 or 60 minutes, and Leica Bond polymer was applied. Leica Bond Mixed Refine (DAB) detection was performed and a Leica hematoxylin counterstain was added. Slides were cleared with xylene, mounted, and scanned in brightfield (20x) using the Aperio ScanScope AT slide scanner. Digital images were imported into Aperio eSlide manager.

### Halo analysis

Images were analyzed using HALO (Indica Labs). Lung tissue was annotated, and the total number of cells was identified using cell by cell analysis. Cells were determined based on thresholds set for nuclear size, segmentation, contrast threshold, and maximum cytoplasm radius. CD8^+^ or CD3^+^ cells were identified by positive staining set by the minimum optical density (OD) above the background (set on regions of non-specific staining). MSLN expression analysis was performed by annotating the tumor region and using the Area Quant analysis to determine the percentage of area with positive MSLN staining set by the minimum OD above the background.

### Statistics

The Student’s *t* test was used to compare normally distributed two-group data. A one-way Anova with post-hoc analysis pairwise for multiple comparisons was used to compare data from experiments with more than two groups. Survival curve analysis was performed using the Log-rank (Mantel-Cox) and Gehan-Breslow-Wilcoxon tests. All error bars represent standard deviation (SD).

## Acknowledgements

The authors thank Matthias Stephan for the ID8_VEGF_ cell line; Amanda Koehne for pathology assistance; Sunni Farley, Jeffrey Williams, and Savanh Chanthaphavong for histopathology assistance; Edison Chiu, Nicolas Garcia and Aesha Vakil for assistance with experiments; the Fred Hutchinson Cancer Research Center Flow Cytometry Core for technical support; Deborah Banker for critical review of the manuscript; and all members of the Greenberg lab for thoughtful and critical discussion.

## Figure Legends

**Supplemental Figure 1. Fas Ligand knockout validation in ID8**_**VEGF**_ **tumor cells**.

Sequence verification of FasL insertion/deletion (indel) in CRISPR/Cas9 targeted ID8_VEGF_ tumor cells. The E9 clone contains a single insertion event in Exon 1 of the FasL gene (indicated by red arrow). Sequencing analysis performed using Synthego’s Inference of CRISPR Edits (ICE) software tool.

## References

1. Lheureux, S., Karakasis, K., Kohn, E. C. & Oza, A. M. Ovarian cancer treatment: The end of empiricism? Cancer 121, 3203–3211 (2015).

2. Vaughan, S., Coward, J. I., Bast, R. C., Berchuck, A., Berek, J. S., Brenton, J. D., Coukos, G., Crum, C. C., Drapkin, R., Etemadmoghadam, D., Friedlander, M., Gabra, H., Kaye, S. B., Lord, C. J., Lengyel, E., Levine, D. A., McNeish, I. A., Menon, U., Mills, G. B., Nephew, K. P., Oza, A. M., Sood, A. K., Stronach, E. A., Walczak, H., Bowtell, D. D. & Balkwill, F. R. Rethinking ovarian cancer: recommendations for improving outcomes. in 11, 719–725 (2011).

3. Gubbels, J. A. A., Belisle, J., Onda, M., Rancourt, C., Migneault, M., Ho, M., Bera, T. K., Connor, J., Sathyanarayana, B. K., Lee, B., Pastan, I. & Patankar, M. S. Mesothelin-MUC16 binding is a high affinity, N-glycan dependent interaction that facilitates peritoneal metastasis of ovarian tumors. Molecular cancer 5, 50 (2006).

4. Chang, K. & Pastan, I. Molecular cloning of mesothelin, a differentiation antigen present on mesothelium, mesotheliomas, and ovarian cancers. Proceedings of the National Academy of Sciences 93, 136–140 (1996).

5. Chang, K., Pastan, I. & Willingham, M. C. Isolation and characterization of a monoclonal antibody, K1, reactive with ovarian cancers and normal mesothelium. International Journal of Cancer 50, 373–381 (1992).

6. Cheever, M. A., Allison, J. P., Ferris, A. S., Finn, O. J., Hastings, B. M., Hecht, T. T., Mellman, I., Prindiville, S. A., Viner, J. L., Weiner, L. M. & Matrisian, L. M. The Prioritization of Cancer Antigens: A National Cancer Institute Pilot Project for the Acceleration of Translational Research. Clinical Cancer Research 15, 5323–5337 (2009).

7. Hassan, R., Thomas, A., Alewine, C., Le, D. T., Jaffee, E. M. & Pastan, I. Mesothelin Immunotherapy for Cancer: Ready for Prime Time? Journal of clinical oncology : official journal of the American Society of Clinical Oncology 34, 4171–4179 (2016).

8. Le, D. T., Wang-Gillam, A., Picozzi, V., Greten, T. F., Crocenzi, T., Springett, G., Morse, M., Zeh, H., Cohen, D., Fine, R. L., Onners, B., Uram, J. N., Laheru, D. A., Lutz, E. R., Solt, S., Murphy, A. L., Skoble, J., Lemmens, E., Grous, J., Dubensky, T., Brockstedt, D. G. & Jaffee, E. M. Safety and survival with GVAX pancreas prime and Listeria Monocytogenes-expressing mesothelin (CRS-207) boost vaccines for metastatic pancreatic cancer. Journal of clinical oncology : official journal of the American Society of Clinical Oncology 33, 1325–1333 (2015).

9. Beatty, G. L., Haas, A. R., Maus, M. V., Torigian, D. A., Soulen, M. C., Plesa, G., Chew, A., Zhao, Y., Levine, B. L., Albelda, S. M., Kalos, M. & June, C. H. Mesothelin-specific chimeric antigen receptor mRNA-engineered T cells induce anti-tumor activity in solid malignancies. Cancer Immunology Research 2, 112–120 (2014).

10. Anderson, K. G., Voillet, V., Bates, B. M., Chiu, E. Y., Burnett, M. G., Garcia, N. M., Oda, S. K., Morse, C. B., Stromnes, I. M., Drescher, C. W., Gottardo, R. & Greenberg, P. D. Engineered Adoptive T-cell Therapy Prolongs Survival in a Preclinical Model of Advanced-Stage Ovarian Cancer. Cancer Immunology Research 7, 1412–1425 (2019).

11. Peter, M. E., Hadji, A., Murmann, A. E., Brockway, S., Putzbach, W., Pattanayak, A. & Ceppi, P. The role of CD95 and CD95 ligand in cancer. Cell death and differentiation 22, 549–559 (2015).

12. Motz, G. T., Santoro, S. P., Wang, L.-P., Garrabrant, T., Lastra, R. R., Hagemann, I. S., Lal, P., Feldman, M. D., Benencia, F. & Coukos, G. Tumor endothelium FasL establishes a selective immune barrier promoting tolerance in tumors. Nature Medicine 20, 607–615 (2014).

13. Oda, S. K., Anderson, K. G., Ravikumar, P., Bonson, P., Garcia, N. M., Jenkins, C. M., Zhuang, S., Daman, A. W., Chiu, E. Y., Bates, B. M. & Greenberg, P. D. A Fas-4-1BB fusion protein converts a death to a pro-survival signal and enhances T cell therapy. The Journal of Experimental Medicine 217, (2020).

14. Heath, R. M., Jayne, D. G., O’Leary, R., Morrison, E. E. & Guillou, P. J. Tumour-induced apoptosis in human mesothelial cells: a mechanism of peritoneal invasion by Fas Ligand/Fas interaction. British journal of cancer 90, 1437–1442 (2004).

15. Chen, L., Park, S.-M., Tumanov, A. V., Hau, A., Sawada, K., Feig, C., Turner, J. R., Fu, Y.- X., Romero, I. L., Lengyel, E. & Peter, M. E. CD95 promotes tumour growth. Nature 465, 492–496 (2010).

16. Yamamoto, T. N., Lee, P.-H., Vodnala, S. K., Gurusamy, D., Kishton, R. J., Yu, Z., Eidizadeh, A., Eil, R., Fioravanti, J., Gattinoni, L., Kochenderfer, J. N., Fry, T. J., Aksoy, B. A., Hammerbacher, J. E., Cruz, A. C., Siegel, R. M., Restifo, N. P. & Klebanoff, C. A. T cells genetically engineered to overcome death signaling enhance adoptive cancer immunotherapy. The Journal of clinical investigation 129, 1551–1565 (2019).

17. Strasser, A., Jost, P. J. & Nagata, S. The Many Roles of FAS Receptor Signaling in the Immune System. Immunity 30, 180–192 (2009).

18. Kägi, D., Vignaux, F., Ledermann, B., Bürki, K., Depraetere, V., Nagata, S., Hengartner, H. & Golstein, P. Fas and perforin pathways as major mechanisms of T cell-mediated cytotoxicity. Science 265, 528–530 (1994).

19. Stromnes, I. M., Schmitt, T. M., Hulbert, A., Brockenbrough, J. S., Nguyen, H. N., Cuevas, C., Dotson, A. M., Tan, X., Hotes, J. L., Greenberg, P. D. & Hingorani, S. R. T Cells Engineered against a Native Antigen Can Surmount Immunologic and Physical Barriers to Treat Pancreatic Ductal Adenocarcinoma. Cancer Cell 28, 638–652 (2015).

20. Snell, L. M., Lin, G. H. Y., McPherson, A. J., Moraes, T. J. & Watts, T. H. T-cell intrinsic effects of GITR and 4-1BB during viral infection and cancer immunotherapy. Immunol Rev 244, 197–217 (2011).

21. Lee, H.-W., Park, S.-J., Choi, B. K., Kim, H. H., Nam, K.-O. & Kwon, B. S. 4-1BB Promotes the Survival of CD8+ T Lymphocytes by Increasing Expression of Bcl-xL and Bfl-1. J Immunol 169, 4882–4888 (2002).

22. Bartkowiak, T. & Curran, M. A. 4-1BB Agonists: Multi-Potent Potentiators of Tumor Immunity. Frontiers Oncol 5, 117 (2015).

23. Zhang, L., Yang, N., Garcia, J.-R. C., Mohamed, A., Benencia, F., Rubin, S. C., Allman, D. & Coukos, G. Generation of a syngeneic mouse model to study the effects of vascular endothelial growth factor in ovarian carcinoma. The American journal of pathology 161, 2295–2309 (2002).

24. Janát-Amsbury, M. M., Yockman, J. W., Anderson, M. L., Kieback, D. G. & Kim, S. W. Comparison of ID8 MOSE and VEGF-modified ID8 cell lines in an immunocompetent animal model for human ovarian cancer. Anticancer research 26, 2785–2789 (2006).

25. Pircher, H., Michalopoulos, E. E., Iwamoto, A., Ohashi, P. S., Baenziger, J., Hengartner, H., Zinkernagel, R. M. & Mak, T. W. Molecular analysis of the antigen receptor of virus-specific cytotoxic T cells and identification of a new V alpha family. European journal of immunology 17, 1843–1846 (1987).

