## Supplementary figures and images for "Engineering adoptive T cell therapy to co-opt Fas ligand-mediated death signaling in ovarian cancer enhances therapeutic efficacy"

### Supplemental Figure 1

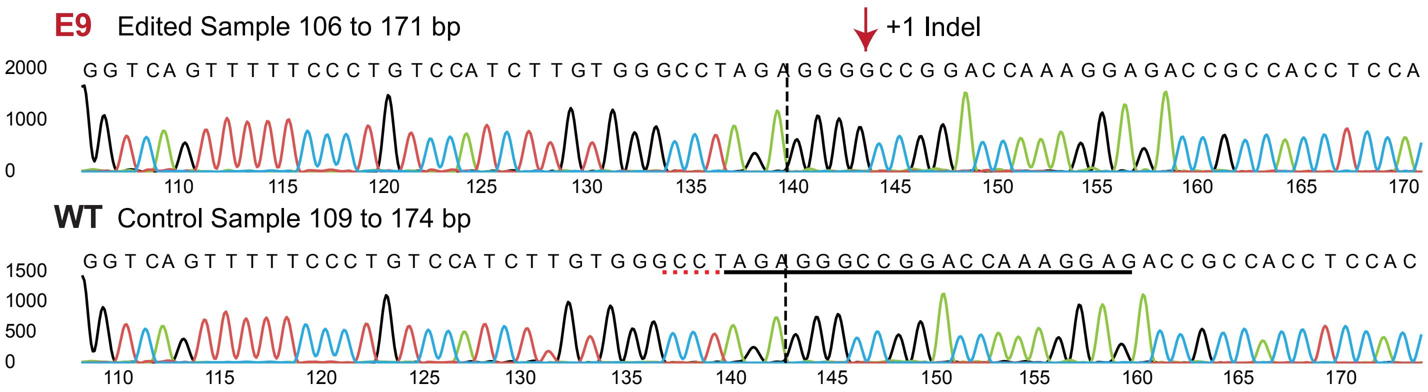
